# Brain microvascular endothelial cell dysfunction during *Plasmodium* infection does not result from astrocyte-derived wingless-related integration site (Wnt)7a deficiency

**DOI:** 10.64898/2026.10.18.760006

**Authors:** Marshall Roedel, Nia Brooks, Jared N. Andersen, Douglas H. Cornwall, Rohan Doddavaram, Kennedy Chastang, Leah Rubner, Tao Liu, Tracey J. Lamb

**Author notes:** **Author contact details:** Marshall Roedel, Nia Brooks, Jared Andersen, Douglas H. Cornwall, Rohan Doddavaram, Kennedy Chastang, Leah Rubner, Tao Liu. Correspondence to: Tracey J. Lamb, 15 N Medical Drive, Room 4280B, Salt Lake City, UT 84112, USA.

## Abstract

Breakdown of the blood brain barrier is a key event leading to vasogenic edema that underpins pediatric cerebral malaria. Brain microvascular endothelial cells (BMECs) downregulate junction proteins in response to *Plasmodium*-infected red blood cells (iRBCs). Very little is known about how glial cell activation in *Plasmodium* infection dysregulates astrocyte-derived steady state signals that control BMEC junction expression. Astrocytes from mice infected with *Plasmodium berghei* ANKA (*Pb*A) exhibit transcriptional changes dominated by interferon signaling pathways. Downregulation of wingless-related integration site (Wnt)7a transcription in astrocytes correlated with transcriptional changes in BMECs indicative of reduced *β*-catenin activity. Incubation of the human astrocyte-like glioma cell-line CCF-STTG1 with interferon-γ (IFN-γ) decreased the general production of Wnt7a in response to *P. falciparum* iRBCs. However astrocyte conditioned medium incubated with IFN-γ strengthened the barrier function *in vitro* when added onto monolayers the human cerebral microvascular endothelial cell line D3 (hCMEC/D3). This data suggests that the response of astrocytes during *Plasmodium* infection may be protective but unable to overcome dysfunctional responses of BMECs to *Plasmodium* iRBCs. To test this hypothesis we overexpressed Wnt7a in the brain tissue of mice using an AAV construct but this did not impart increased survival to the neurological effects of *Pb*A infection. Therefore astrocyte responses during *Plasmodium* infection may not underlie the downregulation of junction proteins on BMECs leading to edema in cerebral malaria.

## Introduction

Malaria, the disease that results from *Plasmodium* infection, remains a serious global health problem resulting in over 600,000 deaths in 2024 (1). One of the leading causes of death from *Plasmodium falciparum* infection is cerebral malaria (CM), a condition that arises when *P. falciparum* infected red blood cells (iRBCs) sequester on the brain endothelium leading to vascular activation, vascular leak and vasogenic edema. This in turn results in loss of consciousness, seizures and coma (2). Most individuals developing CM are children under 5 years old and even with treatment, the condition is still lethal in 20% of patients (3). Those that survive CM are often afflicted with long-lasting disabilities, such as blindness, deafness, epilepsy, motor-disorders, and cognitive and mood disorders (4, 5).

Whilst the cause of vascular leak is multifactorial, downregulation of junction proteins on brain microvascular endothelial cells (BMECs) accompanies vasogenic edema that leads to herniation of the brain and death (6–8). The neurovascular unit (NVU) includes glial cells that function to support blood brain barrier (BBB) integrity (9, 10). Both astrocytes (11) and microglial cells (12, 13) have been shown to become activated during *Plasmodium* infection. This leads to the possibility that breakdown of the BBB may result from lack of support to maintain normal endothelial cell functions related to barrier integrity, rather than endothelial cell activation alone.

BBB breakdown in *Plasmodium* infection can be modelled using the experimental cerebral malaria (ECM) model of *Plasmodium berghei* ANKA (*Pb*A) infection of C57BL/6J mice and leads to death of mice 6 to 10 days post-infection. CD8 T cells accumulate in the brain during pediatric CM (14) and are required for vascular leak in ECM (15, 16). Granzyme B and perforin appear to be central mediators of vascular dysfunction in ECM (17, 18) despite the lack of significant BMEC apoptosis (19). However, there is a prominent downregulation of the tight junction proteins claudin-5 and occludin which normally confer barrier properties of BMECs by preventing para-cellular leak with areas of downregulation corresponding to areas of vascular leak (20).

Astrocytes can support BMEC functions such as regulation of junction proteins to control passage of solutes across the BBB (21, 22). In steady state astrocytes secrete factors that can promote tight junction expression in BMECs. These include the wingless-related integration site (Wnt) ligands Wnt7a and Wnt7b (23–25), Sonic Hedgehog (26, 27), Apolipoprotein E (Apo-E) (28, 29), brain derived neurotrophic factor (BDNF) (30), Cystatin-C (31), and angiotensinogen (32). Astrocytes can respond to inflammation and injury by changing their transcriptional profile as well as their conformation (33, 34). This coincides with an upregulation of the expression of the intermediate filament protein Glial Fibrillary Acidic Protein (GFAP) (35), as observed in pediatric CM (36). These responses can modulate the ability of astrocytes to support the BBB whic may, in turn, disrupt barrier function.

The *Pb*A ECM model has demonstrated a key role of interferon-γ (IFN-γ) in the breakdown of the BBB during *Plasmodium* infection (37), possibly by promoting the accumulation of CD8 T cells in the CNS (16). IFN-γ can act directly on BMECs, facilitating cross-presentation of *Pb*A-derived antigens by BMECs mediating recognition of *Plasmodium*-derived epitopes by CD8 T cells (38). The indirect effects of IFN-γ are less well described. Here we have used the *Pb*A model of ECM in combination with *in vitro* cultures of human astrocyte-like cells exposed to *P. falciparum* iRBCs to determine that astrocytes display a strong interferon-mediated transcriptional response that is associated with downregulation of Wnt7A, but may not be the underlying cause of BBB disruption. Rather astrocytes appear to upregulate barrier promoting molecules in response to *Plasmodium* iRBCs. These data suggest that a failure of barrier promoting activities by activated astrocytes in *Plasmodium* infections is not the primary driver of junction protein downregulation on BMECs.

## Materials & methods

### Ethics statement

This study was approved by the University of Utah Institutional Animal Care Use Committee (Protocol Number #2051). All procedures were carried out in accordance with institutional guidelines for the care and use of animals.

### Rodent *Plasmodium* infections

Female C57BL/6J mice aged 4-8 weeks were purchased from The Jackson Laboratory (line #000664; Bar Harbor, Maine, USA) and were housed under standard conditions. All mice had access to food and water *ad libitum. Pb*A infections were initiated intraperitoneally with 0.5 x 10^6^ iRBCs. Peripheral parasitemia was quantified in 300-500 red blood cells (RBCs) on thin blood smears stained with Hema 3 Stat Pack (Fisher Scientific). Anemia was quantified using a Beckman Coulter Z2 coulter counter on blood diluted 1:50,000 in Isoton solution. Cachexia was monitored by weighing mice on a top pan balance. Clinical scoring was conducted according to the protocol previously described (39). Briefly the 20 behavioral parameters were scored between a scale of 2 (healthy) to 0 (unhealthy) and then summed. Scores below 15 are considered as displaying neurological symptoms of *Pb*A infection.

### Whole brain nuclei isolation for single nuclear RNA (snRNA) sequencing

Brains were harvested and aseptically diced into 1 mm^3^ cubes with a scalpel. Brain tissue from each mouse was digested for 30 minutes in 5 ml of Earle’s Balanced Salt Solution (EBSS) with Calcium and Magnesium (Gibco #24010) and 100 U of papain (Worthington # LK003178) in a gentleMACS Octo Dissociator with Heaters using program 37C_ABDK_01. Miltenyi magnetic separation beads were used to remove myelin debris according to the manufacturer’s instructions (Miltenyi #130-096-433). Nuceli were extracted by lysing cells in a solution 0.1% NP-40 in Hanks Balanced Salt Solution (HBSS) on ice for 3 minutes. After centrifguation, the supernatant was decanted and the pellet was resuspended in HBSS with 1:1000 dilution of 7-amino-actinomycin D (7-AAD) (ThermoFisher #06993-50). The nuclei were sorted on a Miltneyi Tyto cell sorter. Single nuclear libraries of 10,000 cells per condition were generated by the University of Utah DNA Sequencing Core using 10X Chromium Next GEM Single Cell 5’ Version 2 (PN-1000263). Sequencing was performed with an Illumina Miseq with a target of 20,000 reads per cell.

### Mouse brain endothelial cell isolation for single cell RNA (scRNA) sequencing

Brains were harvested and aseptically diced 1 mm^3^ cubes with a scalpel. Brain tissue from each mouse was digested for 30 minutes using the Miltenyi gentlMACS system in 5 ml of EBSS with calcium and magnesium (Gibco 24010) and 100 U of papain (Worthington # LK003178) in a gentleMACS Octo Dissociator with Heaters and program 37C_ABDK_01. Miltenyi magnetic separation beads were used to remove myelin debris (Miltenyi #130-096-433) and CD45+ cells negatively selected according to the manufacturer’s instructions (Miltenyi #130-052-301). Beads were then used to isolate the remaining CD31+ cells (Miltenyi # 130-097-418). Single cell libraries of 10,000 cells per condition were generated by the University of Utah DNA Sequencing Core using 10X Chromium Next GEM Single Cell 5’ Version 2 (PN-1000263). Sequencing was performed with an Illumina Miseq with a target of 20,000 reads per cell.

### SnRNA and scRNA sequencing and data processing

Analysis of the single nuclei and cell dataset were completed using Seurat (40). For quality control of the whole brain snRNA sequencing dataset, nuclei with a mitochondrial transcript percentage of above 2.5% or with fewer than 400 features were excluded from the analysis. For quality control of the scRNA sequencing dataset, CD31+ cells with a mitochondrial transcript percentage of above 10% a feature number outside of the range of 300-5000 were excluded. The two conditions from each experiment were merged and then scaled and normalized using the SCTransform functions to find 5000 variable features. A principal components analysis (PCA) was performed and the two conditions in each experiment were integrated using a canonical correlation analysis (CCA). For the clustering analysis 50 dimensions were used at a resolution of 0.1 for the snRNA sequencing dataset and 0.2 for the scRNA sequencing dataset. Cluster identities were determined using established markers and the commands FeaturePlot, VlnPlot, and FindMarkers were used for differential gene expression analysis. Standard pipelines for the “EnhancedVolcano” package and the and “clusterProfiler” package were used to generate volcano plots and Gene Ontology (GO) enrichment plots respectively.

### Culture of human Plasmodium falciparum

De-identified O positive human RBCs were obtained from the the University of Utah blood bank. Prior to infection with *Plasmodium,* RBCs were washed RPMI 1640 (Thermofisher #11875093) three times. *P. falciparum* cell lines 3D7 and DD2 (a kind gift from Paul Sigala, University of Utah) were cultivated in RBCs at 2-4% hematocrit in RPMI 1640 media containing 5g/L of albumax (Thermofisher #11020021), 30mg/L hypoxanthine (Sigma Aldrich #H9377-1G), 4g/L of glucose, and 20 mM of sodium pyruvate (Thermo Fishter # 11360070). *Plasmodium* cultures were maintained in 90% nitrogen, 3% oxygen, and 7% carbon dioxide at 37°C.

### Astrocyte responses to *Plasmodium*

The CCF-STTG1 cell line (ATCC #CRL-1718) was grown to confluence in RPMI in 10% FBS (Gibco #11875093). Cells were incubated with human naïve RBCs, iRBCs (*P. falciparum* 3D7 or DD2 schizonts), or from lysates from naïve or iRBCs at a ratio of 10 RBCs / astrocyte. Schizonts were isolated from *P. falciparum* cultures through magnetic isolation using Miltenyi LS magnetic isolation columns (Miltenyi #130-042-401) and QuadroMACS magnetic separators (Miltenyi #130-091-051)(41). Recombinant IFN-*γ* (Peprotech #300-02) or TNF-α (Preprotech #300-01A) was added at a concentration of 10 ng/ml. After 24 hours the supernatant was frozen at -20°C until used. Wnt7A secretion in the supernatant was quantified by ELISA (Aviva # OKEH01755) according the manufacturer’s instructions. EphA2 and platelet derived growth factor-D (PDGFD) were quantified in the supernatant using Luminex technology (R and D systems).

### Confocal microscopy of brain sections

Brains were bisected and incubated in 4% formaldehyde at room temperature for one hour prior to sequential increasing sucrose concentrations of 15% and 30% at 4°C for 24 hours per concentration. Brains were cryopreserved in Optimal Cutting Temperature (OCT) (Electron Microscopy Sciences #62550-01) and stored at -80°C. 10µm thick cryosections were rehydrated in PBS for 10 minutes, and antigen retrieval performed by incubation in 1% sodium dodecyl sulfate (SDS) solution for five minutes. Sections were blocked with 1% BSA in a 0.2% Triton-X 100 solution for 1 hour at room temperature before being incubated with antibody overnight at 4°C. After washings sections were mounted with Invitrogen Fluoromount-G containing DAPI to visualize nuclei (Invitrogen # 50-112-8966).

### Wnt7a - expressing Adeno-associated virus (AAV) infection of mice

C57BL/6J mice were intravenously injected with 2.1x10^11^ viral genomes (vg) of either the Wnt7a ligand AAV-PHP.eb-WNt7aK190A-p2A-GFP or GFP control AAV-PHP.eB-GFP (a kind gift from Benoit Vanhollebeke, University of Brussels) as previously published (42). Two weeks after AAV injection brain GFP expression was quantified through immunofluorescence on brain sections after incubation with a 1:500 dilution of GFP booster (Chromotek #gba488) or were infected with *Pb*A to quantify the pathogenesis of infection.

### Depletion of microglia

C57BL/6J mice aged 5-7 weeks were gavaged daily with 600 µg of Pexidartinib (PLX-3397: MedChem Express #HY-16749) or PLX-5622 (MedChem Express #HY-114153) dissolved in 100 µl of olive oil from 7 days before the infection with *Pb*A until death. Microglia depletion was verified with confocal microscopy of brain sections incubated with 1:200 dilution of anti-CD11b BV605 (Biolegend #101257).

### Generation of astrocyte conditioned media

CCF-STTG1 astrocyte-like cells were incubated for 24 hours with lysed naive RBCs or *P. falciparum* 3D7 schizont lysates at a ratio of 10 RBCs per astrocytes in the presence or absence of 10 ng/ml of recombinant IFN-*γ* (Peprotech #300-02).

### Electric Cell-substrate Impedance Sensing (ECIS) analysis of brain microvascular endothelial cell barrier function

Barrier function of human brain endothelial cell lines was assessed using the Xcelligence RTCA instrument (43). 7.5 X 10^4^ human cerebral microvascular endothelial cells (hCMEC/D3) (44) (SigmaAldrich #SCC066) were seeded into each well of an Agilent 96-well E-Plate (Agilent # 5232368001) and incubated in an xCELLigence RTCA W380 Real-Time Cell Analyzer W / 1X96 (TT Instruments # 8171 4D04) for 59 hours at 37°C with 5% CO_2_ in 100 *μ*l of EndoGro-MV complete media (SigmaAldrich #SCME004). At 59 hours each well received fresh media. At 60 hours, 70 *μ*l of media from each well was removed and replaced with an equal volume of astrocyte conditioned media. Impedance was quantified every minute for 100 measurements, then every 3 minutes for the next 200 measurements, and every 15 minutes for the remaining measurements to cover a total period of 24 hours. Cell index, the measure of barrier function, was calculated by subtracting the media only baseline impedance from the measured impedance. To compare conditions the cell index at the time point when the conditioned media was added was subtracted from the experimental time point as previously described (45).

### Statistical Analysis

Statistical tests were carried our using Prism software (Graphpad) and Rstudio software. Log-rank Mantel-Cox tests were applied to test for significant differences in survival analyses. Comparisons between two groups were carried out using a two-tailed Mann-Whitney U tests. More than two groups were compared using nonparametric Kruskal-Wallis tests followed by a Dunn’s multiple comparison test or using General Linear Modelling (GLM) on models with normally destributed residuals (Minitab Inc) with P values reported on minimal models that included first order interactions. p<0.05 was considered statistically significant all cases.

### Data availability statement

The single nuclear and single cell RNA sequencing data sets described in this study can be accessed via the NCBI Gene Expression Omnibus data set GSE348481.

## Results

### Activated astrocytes upregulate interferon signaling upon *Pb*A infection

To quantify how astrocytes respond during *Pb*A infection, we undertook snRNA sequencing from whole brain at day 6 post infection to quantify the transcriptional pathways activated in astrocytes. Cluster analysis from the brain-derived nuclei yielded 16 clusters (**Fig. 1A**). Cluster identities were determined using the top markers from each cluster. Clusters 6 and 14 expressed high module scores for astrocytes based on expression of the astrocyte markers *Aldh1l1, Gfap*, *Aqp4*, *Slc1a3*, *Atp1b2*, and *S100b* (**Fig. 1B**) (46–48). High expression of the gene encoding Paired Box 3 (Pax 3) in cluster 14 was indicative that this cluster contained Bergmann glia, specialized astrocytes restricted to the cerebellum (49, 50). Compared to the nuclei isolated from the brains of naïve C57BL/6J mice, 1764 genes were differentially expressed in mature astrocytes in cluster 6 in mice infected with *Pb*A, 1167 of which were upregulated (**Fig. 1C**). Of the 339 differntially expressed genes in the Bergmann glia cluster, 206 were upregulated (**Fig. S1A).**

**Figure 1:**
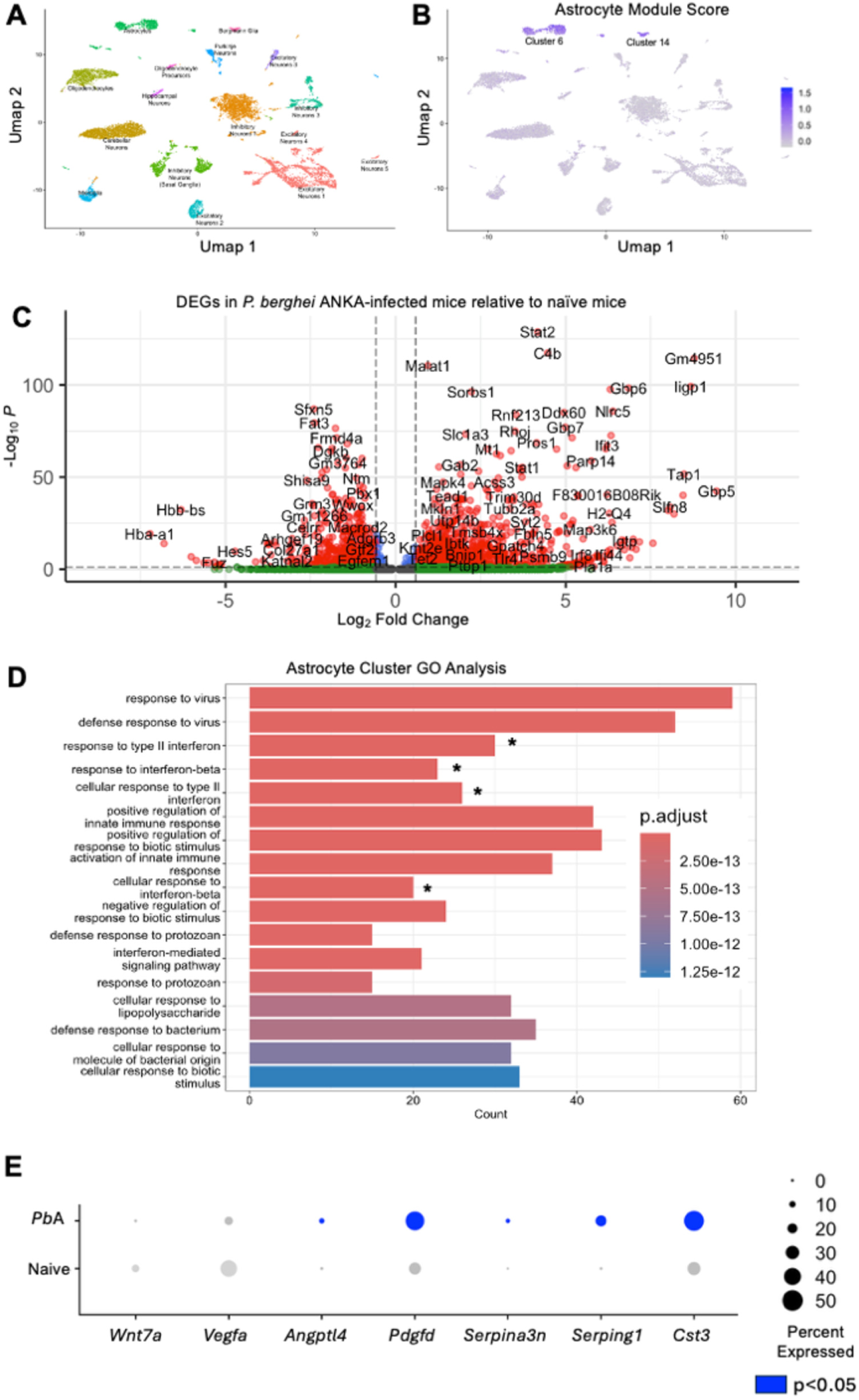
Astrocytes upregulate interferon response genes and alter BBB regulation genes in response to *Pb*A infection. Single nucleus RNA-sequencing (snRNA seq) clustering analysis of whole brains of 2 pooled naïve control mice and 2 mice infected with *Pb*A yields 16 clusters (**A**). Two clusters highly express astrocyte markers (*Slc1a3*, *Atp1b2*, *Gfap*, *Aldh1l1*, *Aqp4*, and *S100b*), one is general astrocytes, the other expresses Bergmann Glia markers (**B**). Volcano plot of differentially expressed genes in astrcoytes from *Pb*A-infected mice relative to astrocytes from naive mice (**C**). The gene ontology analysis of the 500 most highly upregulated genes in the general astrocyte cluster from *Pb*A-infected mice relative to naive mice (**D**). Terms marked with a * indicate pathways dominated by interferon signaling. A dot plot of genes that regulate blood brain barrier function that are significantly differentially expressed in the astrocytes of *Pb*A-infected and naive mice (**E**). p<0.05 was considered statistically significant.

By inputting the top 500 most highly upregulated genes from cluster 6 (mature astrocytes) (**Fig. 1C**), and 200 from the cluster 14 (Bergmann Glia) (**Fig. S1A**) into a Gene Ontology (GO) analysis (51) we determined that many of the most significantly enriched biological processes centered on interferon signaling. Significant terms included “response to type II interferon”, “response to type I interferon”, and “response to interferon signaling pathway” (**Fig. 1D**; **Fig. S1B**) suggesting that either type II or type I interferon signaling is a major driver of the astrocyte response during *P. berghei* ANKA infection (**Fig. 1D**). This finding is consistent with the previous literature which supports the idea that interferon signaling to be an important driver in ECM (37) (52). It also supports the data demonstrating a significant interferon signaling trasncriptional program in microglia after *Plasmodium* infection (12)( **Fig. S2A**). Although microglia have been shown to play an important role in BBB maintenance (53) and may be important contributors to *Pb*A induced BBB dysfunction through promoting CD8 T cell trafficking to the brain (54) (**Fig. S2C**), depletion of microglia did not protect mice from lethal *Pb*A infection (**Figs. S2D and S2E**).

Many of the differentially expressed genes were those known to be involved in astrocyte regulation of BBB integrity (**Fig. 1E, Fig S1C**). *Wnt7a* (24, 25, 42), an important promoter of BBB function (24, 25, 42), was downregulated in astrocytes from *Pb*A-infected mice compared to naive mice. There was significant upregulation of *Serping1* (55, 56), *Pdgfd* (57, 58) and *Cst3* (31, 59) all of which promote barrier function (24, 25, 31, 42, 55, 57) suggesting that astrocytes increase the secretion of molecules that would othersie maintain BBB integrity. In support of this hypothesis, transcription of *Vegfa* encoding Vascular Endothelial Growth Factor A (VEGFA) which destabilizes the BBB (60) was downregulated and *SerpinA3n* a gene encoding a serine protease inhibitor (61, 62) which inhibits granzyme B in mice (63) was upregulated. This would be predicted to be protective in ECM given the necessity of granzyme B for ECM pathogenesis (17). Collectively the transcriptional results indicate that interferon signaling is a significant influence on astrocyte transcriptional programs in *Pb*A-infected mice and this is associated with concomitant modulation of transcription for genes that modulate BBB integrity.

### Transcriptional changes in BMECs in mice infected with *P. berghei* ANKA reflect modulation of astrocyte proteins that maintain the BBB

To determine how astrocyte-mediated supportive pathways may impact on BMECs in *Pb*A infected mice, we created modules of genes upregulated by signaling by wingless related integration site 7a (Wnt7a), Cystatin C (encoded by *CST3*), platelet-derived growth factor D (PDGFD) and vascular endothelial growth factor A (VEGFA) (**Table 1**) and quantified module activity scores in BMECs isolated from the brain at day 6 post-infection (**Fig. 2A**). Our BMEC single cell data set yielded cells that almost ubiquitously expressed *Pecam1* which encodes the endothelial cell marker CD31, as well as other endothelial markers like Solute carrier 2a1 (Slc2a1) and vascular-endothelial cadherin encoded by *Cdh5* (**Fig. S3B**). Cluster analysis resulted in six clusters (**Fig. 2B**). Clusters 4-6 expressed also markers of artrioles (ART-Cluster 4: *Mgp*, *Bmx*, and *Stmn2* (64), pericytes or vascular smooth muscle cells (MCD-Cluster 5: *Myl9*, *Vtn*, *Atp13a5*, and *Pdgfrb*) or astrocytes (Cluster 6: *Mlc1* and *Slc1a* (65, 66)) (**Fig. S3B**). Given that cluster 6 expressed very low levels of *Pecam1* this cluster was ommitted from further analyses.

**Figure 2:**
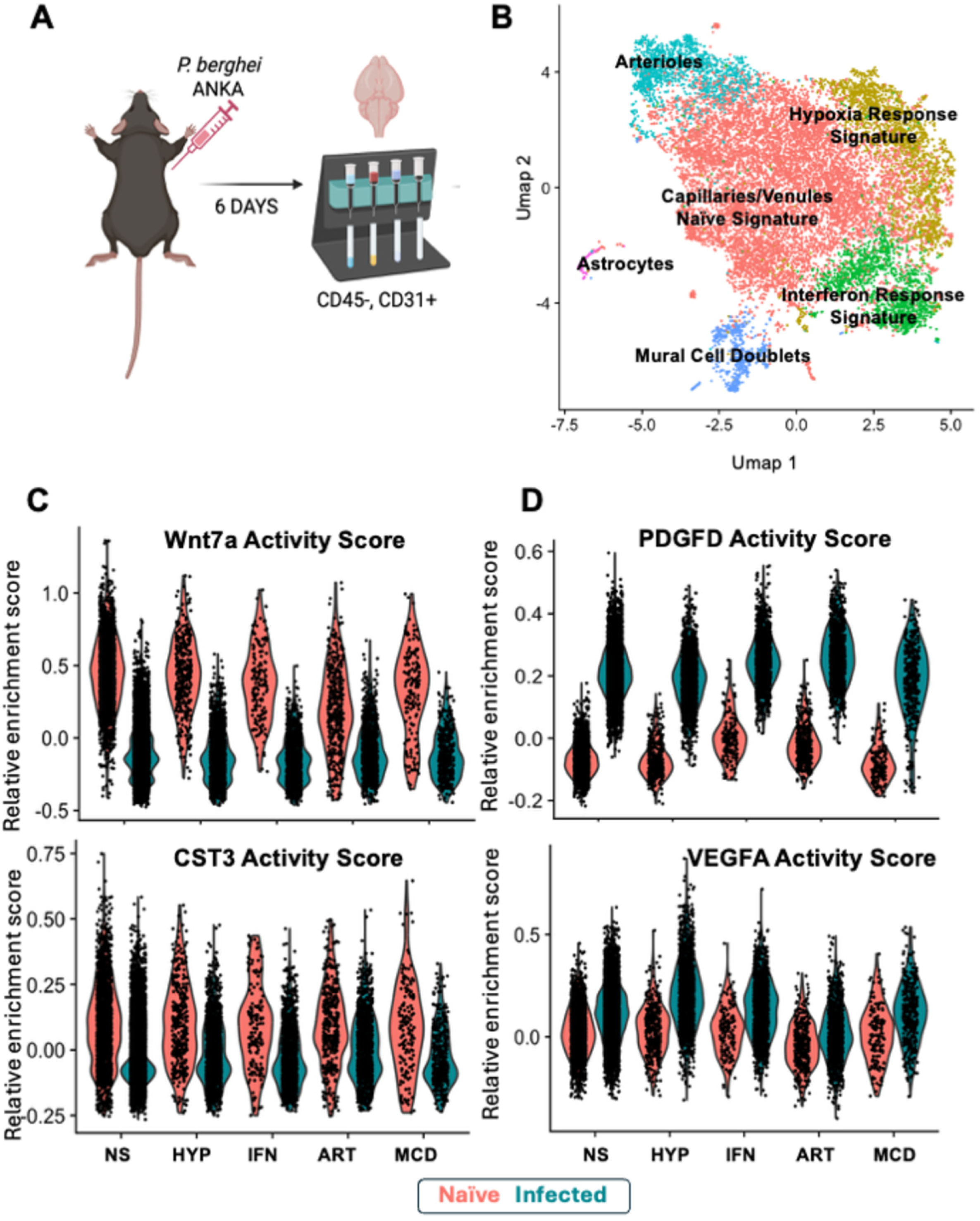
Transcriptional changes in BMECs from *Pb*A-infected mice reflect changes in astrocyte BBB regulation genes. Brain endothelial (CD45-,CD31+) cells were magnetically isolated from 3 naïve and 3 *Pb*A-infected mice for single-cell RNA sequencing (scRNA seq) (**A**). Cluster analysis yields 5 BMEC clusters that represent capillaries and venules with a naïve transcriptional signature (NS), with a hypoxia reactive signature (HYP), or an interferon response signature (IFN) as well as arterioles (ART), and a cluster enriched in mural cell markers suggesting doublets (MCD) (**B**). Module scores representing the transcriptional response to Wnt7a and CST in the different BMEC clusters from *Pb*A-infected mice (**C**). Module scores representing the transcriptional response to Wnt7a and CST in the different BMEC clusters from *Pb*A-infected mice (**D**). p<0.05 was considered statistically significant.

**Table 1:**
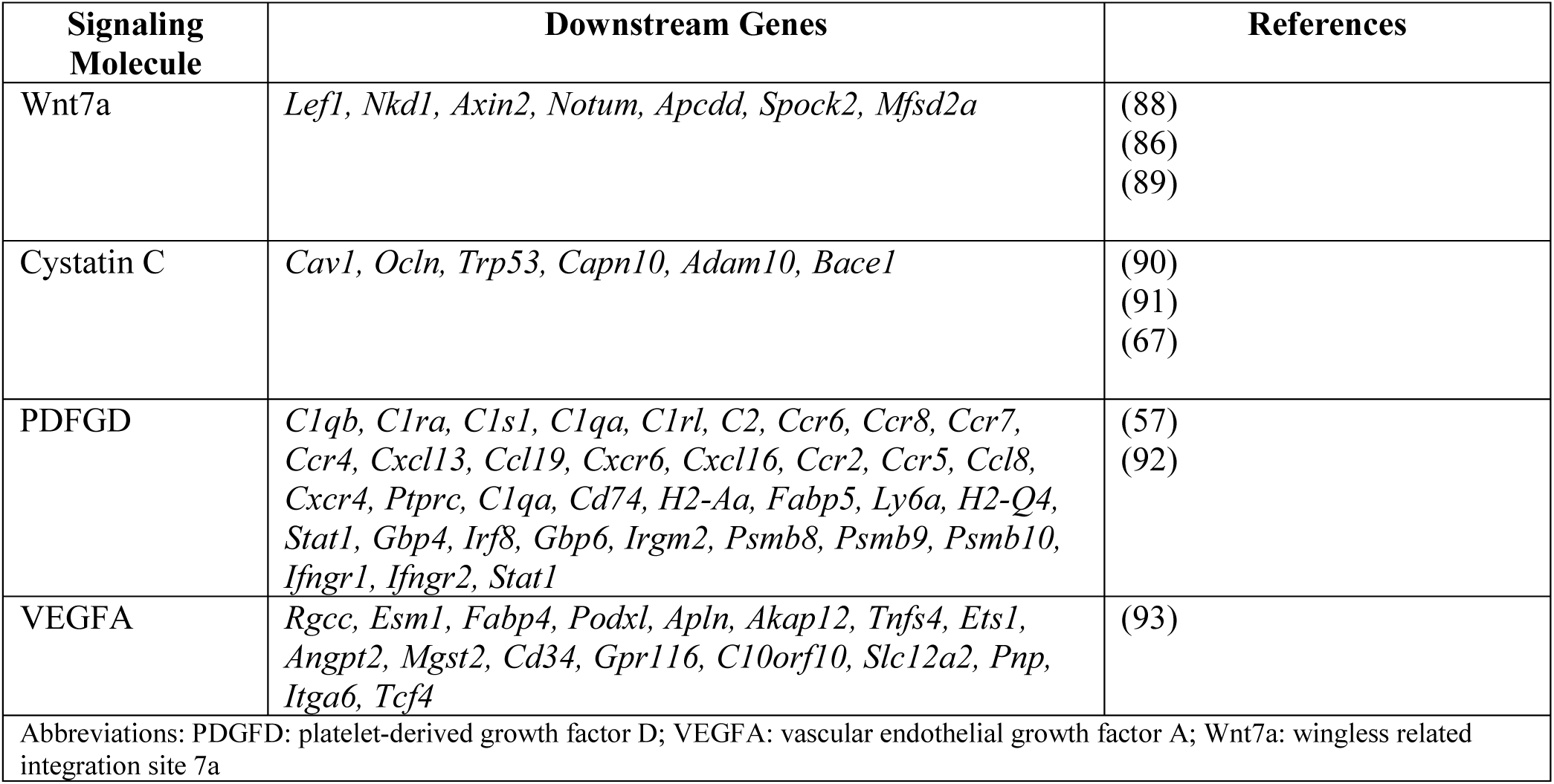
Modules for endothelial response to differentially expressed BBB regulation genes.

Clusters 1-3 were not clearly distinguished by markers associated with cell type, but by those associated with responses to specific stimuli, suggesting that these clusters are composed of capillary and venous endothelial cells in varying states. Relative to clusters 2 and 3, cluster 1 expressed higher levels of *Nkd1* and *Apcdd1* which are activated by *β*-catenin activity, an important pathway for establishing barrier function in brain endothelial cells (67). Along with less differentially expressed genes this suggests that cluster 1 may be more in a naive transcriptional state (NS). The dominant transcriptional sigatures quantified in by GO analysis indicated that cluster 2 is undergoing a ‘response to hypoxia’ and ‘response to decreased oxygen levels’, suggesting this cluster is composed of BMECs in a hypoxic response state (HYP) (**Fig. S3C**). Cluster 3 GO analysis yielded terms including ‘defense response to virus’, ‘activation of innate immune response’, and ‘cellular response to interferon-*β*’, suggesting that these are immune activated endothelial cells with significant interferon signaling pathways activated (IFN) (**Fig. S3D**). Both hypoxia and immune processes have been hypothesized to contribute to pathogenesis in ECM although the interactions between these two processes is not fully understood (68–70).

Consistent with the downregulation of *Wnt7a* transcription in astrocytes from *Pb*A-infected mice, BMECs concomitantly had a reduced Wnt activity score across all 5 clusters of CD31+ cells analyzed (**Fig. 2C**). However the module activity score for signaling in response to Cystatin C also decreased in the *Pb*A-infected mice relative to naïve mice (**Fig. 2C**) which occurred despite upregulation of *Cst3* transcript in astrocytes (**Fig 1E**). On the other hand the module activity score for PDGFD and VEGFA increased in *Pb*A-infected mice, albeit this was more striking for PDGFD (**Fig. 2D**). Therefore BMECs display a pattern of transcriptional changes that include pathways from regulatory proteins produced by astrocytes.

### IFN-γ reduces Wnt7a secretion by astrocyte-like CCF-STTG1 cells

Given the strong IFN signaling pathways initiated in astrocytes during *Pb*A infection and the prominent decreased Wnt7a activity score in the transcriptional pathways in BMECs from *Pb*A-infected mice (**Fig. 2C**), the role of IFN signaling in Wnt7a production from astrocytes was investigated using cultured human the astrocyte-like cells (**Fig. 3A**). Astrocytes were cultured with *P. falciparum* iRBCs in the presence or absence of recombinant human IFN-γ. To verify that astrocytes could respond to IFN-γ, we quantified the effects on the cleavage / secretion of soluble receptor tyrosine kinase EphA2. Increased soluble EphA2 has been previously reported during both human and mouse *Plasmodium* infections (71). IFN-γ induced significant amounts of cleavage / secretion of EphA2 from the cultured astrocytes (GLM IFN-γ p=0.000)(**Fig. 3B**).

**Figure 3:**
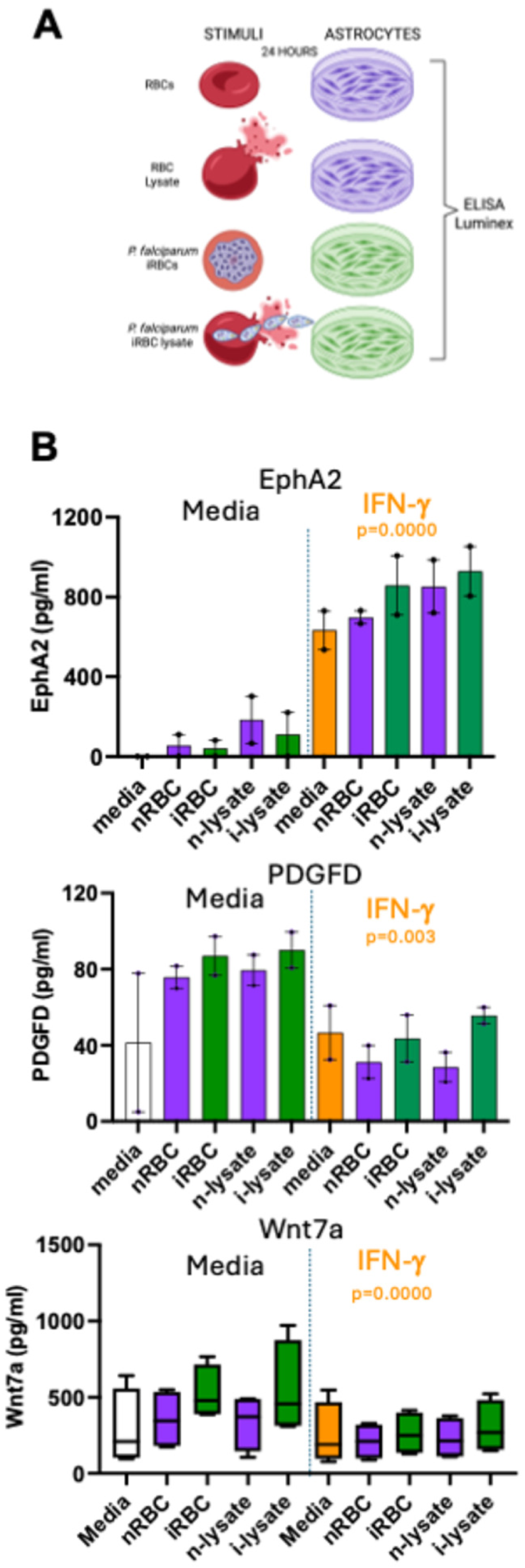
IFN-*γ* reduces Wnt7A secretion in the CCF-STTG1 glioma cell line. The experiment design for assessing Wnt7A secretion by the human glioma cell line CCF-STTG1 in response to exposure to *P. falciparum* iRBCs and recombinant IFN-*γ* (**A**). Secretion of EphA2, PDGFD and Wnt7a was quantified after incubation of cultures for 24 hours (**B**). General Linear Modelling containing 1st order interactions p<0.05 was considered statistically significant. EphA2; ephrin receptor A2; iRBCs: *Pb*A-infected red blood cells; i-lysate: iRBC lysate; nRBC: naive red blood cells; n-lysate: nRBC lysate; PDGFD: platelet derived growth factor D; Wnt7A: wingless-related integration site 7A.

The addition of *P. falciparum* iRBCs increased the production of Wnt7A from astrocytes (GLM, stimulus p=0.000) but this was suppressed when IFN-γ was added to the culture (GLM, IFN-γ p=0.000). This supports the hypothesis that Wnt7a transcript may be suppressed by the strong IFN transcriptional signatures (**Fig. 1D**) and the reduced Wnt7a activity score in BMECs (**Fig. 2C**) observed in astrocytes in *Pb*A infected mice. Interestingly, we found small amounts of PDGFD were produced by CCF-STTG1 cells and that IFN-γ reduced the amount of astrocyte-derived PDGFD (GLM IFN-γ p=0.003). PDGFD is a high affinity agonist for PDGFRβ, a receptor which can promote neurological vascular integrity (57). Whilst PDGFD is normallyassociated with production by BMEC as opposed to astrocytes, there was a significant upregulation in transcription by astrocytes from *Pb*A infected mice relative to naive mice (**Fig. 1E**). This data demonstrates tha *Plasmodium* iRBCs can induce Wnt7A from astrocytes but that this response is significantly dampened by the presence of IFN-γ.

### Infection with AAV constructs that overexpress Wnt7a did not protect mice from death in *P. berghei* ANKA infection

Wnt7a / β-catenin signaling has been shown to be important for the development of the BBB (72), and Wnt7a supplementation been shown to prevent disruption of the BBB in SARS-CoV2 infection (25). To test the hypothesis that suppression of Wnt7a production by astrocytes may underlie fatal disruption of the BBB during *Pb*A infection, mice were injected with an adenovirus construct that expresses an engineered Wnt7a ligand (42) two weeks prior to infection with *Pb*A (**Fig. 4A**). The success of the AAV injection was not significantly different between the mice injected with AAV Wnt7a and the AAV GFP control construct (Mann-Whitney U test, p = 0.4) (**Fig. 4B**). Supplementation of Wnt7a did not improve the survival rate of mice infected with *Pb*A (Kaplan Meier, p = 0.2553) (**Fig. 4C**). Prior to death there was no difference in the severity of neurological manifestations (Mann-Whitney U test, p = 0.3283) (**Fig. 4D**), weight loss (Mann-Whitney U test, p = 0.8714) (**Fig. 4E**) or circulating parasitemia (Mann Whitney-U test, p = 0.0592) at day 5 post-infection (**Fig. 4F**). Collectively, these data suggest that the viral expression of engineered Wnt7a ligand is not sufficient to protect from BBB disruption leading to death from *Pb*A infection.

**Figure 4:**
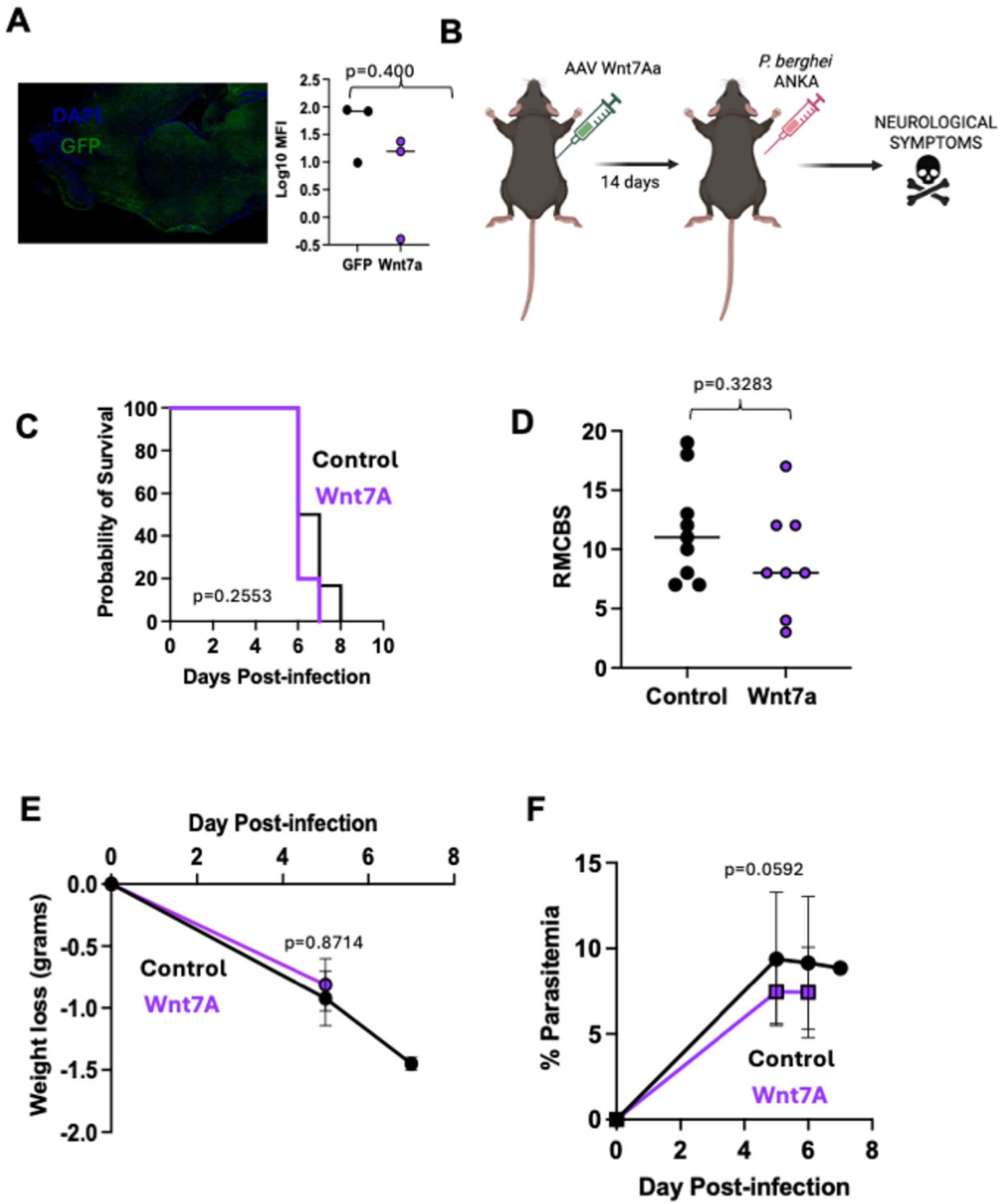
Over expression of engineered Wnt7a ligands does not protect against death in ECM. Mice were intravenously injected with Wnt7a-GFP AAV or the GFP-only AAV construct two weeks before infection with *Pb*A (**A**). The level of construct in the brain was quantified using confocal microscopy of brain tissue at 14 days post injection (**B**). The levels of fluorescence was not different between mice injected with Wnt7a and GFP control constructs (Mann-Whitney U test p=0.04). Survival of mice post infection with *Pb*A was compared between mice previously injected with Wnt7a-GFP AAV (n=8) compared to GFP-only AAV (n=9) (**C**) along with clinical score (**D**), weight (**E**), and parasitemia (**F**). A difference in survival was tested using a Log-rank Mantel-Cox test. Mann-Whitney U test was used to compare differences between groups. p<0.05 was considered statistically significant.

### Conditioned media from astrocytes exposed to *P. falciparum* RBCs improves barrier integrity in cultured hCMEC/D3 endothelial cells

To determine whether IFN-γ signaling on astrocyte responses contributes to the failure of BMECs to maintain the integrity of the BBB, the hCMEC/D3 cell line was incubated with astrocyte conditioned media. Astrocytes were incubated for 24 hours in the presence or absence of recombinant IFN-γ along with lysate from *P. falciparum* iRBCs (**Fig. 5A**). The effects of proteins from lysed RBCs were controlled for by incubation with lysate from naive RBCs. The integrity of hCMEC/D3 monolayers was quantified using ECIS (**Fig. 5B**). The addition of recombinant TNF-α had a temporary but modest effect in disrupting the monolayer as expected (**Fig. 5C**). There was also a small but temporary disruption on the monolayer after the addition of recombinant IFN-γ (**Fig. 5C**). On the other hand the addition of RBC lysate (naive or infected) significantly disrupted the monolayer, with lysed *P. falciparum* iRBCs inducing the most disruption as expected (**Fig. 5C**). When IFN-γ astrocyte conditioned media was added to hCMEC/D3 cell monolayers the monolayers had increased impedance in all conditions (IFN-γ-GLM p=0.029) (**Figs. 5D and 5E**) suggesting that astrocyte-derived soluible molecules led to an improved barrier integrity, despite no increase in the integrity of hCMEC/D3 monolayers when incubated with IFN-γ alone (**Fig. 5C**). This data support the hypothesis that in *Plasmodium* infection, IFN signaling in astrocytes leads to responses that attempt to maintain the BBB in the face of *Plasmodium*-induced inflammation.

**Figure 5:**
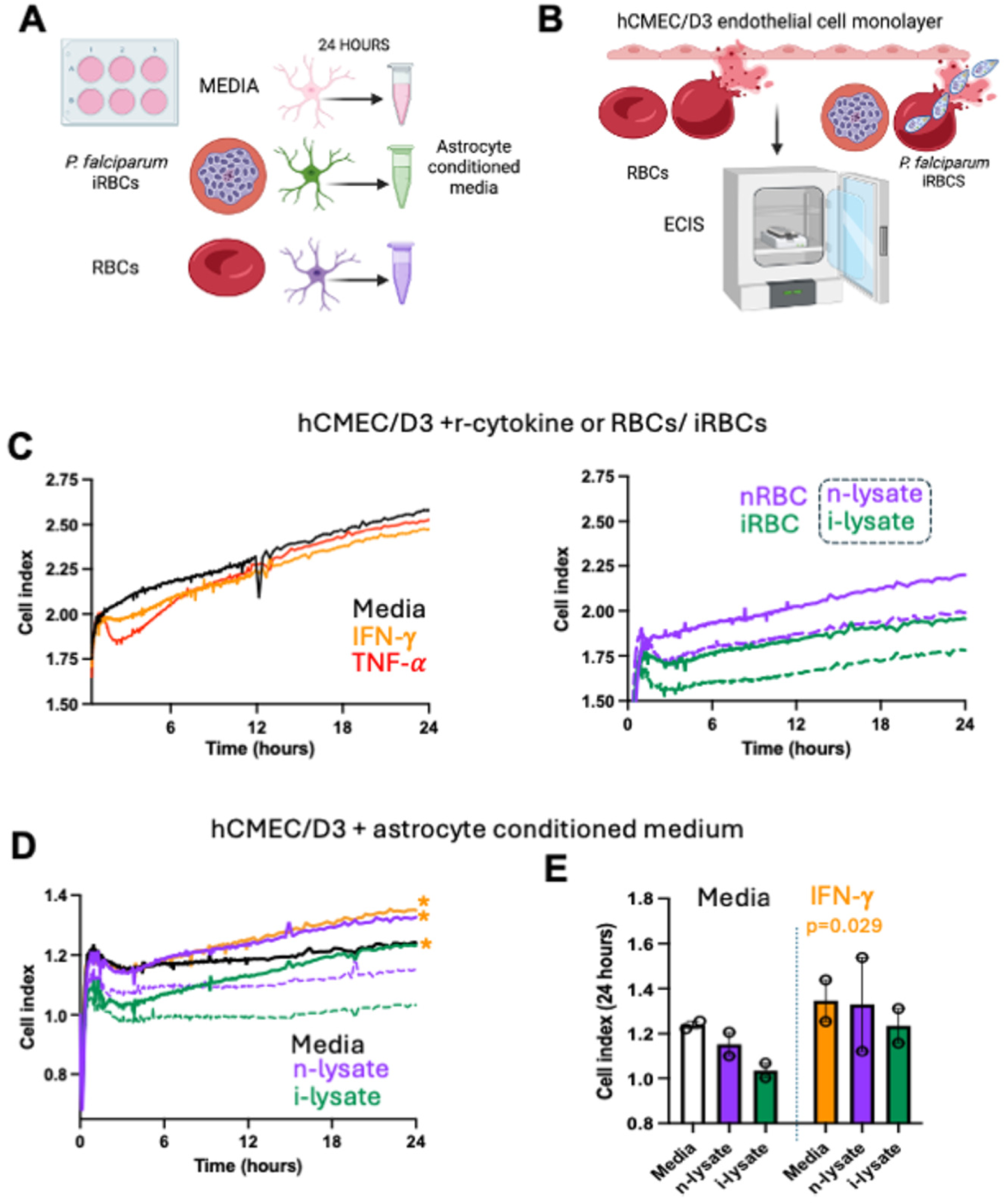
Conditioned media from astroglioma cell line CCF-STTG1 incubated in IFN-*γ* increases barrier properties in human brain endothelial cell line hCMEC/D3. Conditioned media was made by incubating CCF-STTG1 astroglioma cell line with IFN-*γ* and *P. falciparum* iRBC lysate (**A**). Endothelial cell line barrier properties of were measured on hCMEC/D3 monolayers using ECIS assays over 24 hours (**B**). The impedance of hCMEC/D3 monolayers incubated with recombinant cytokines (**C**) or incubation with *P. falciparum* iRBCs for 24 hours. The addition of conditioned medium from astrocytes cultured with RBC lysates in the presence or absence of IFN-γ to hCMEC/D3 monolayers (**D**) and the cell index at 24 hours (**E**). Lines marked with * indicates stimuli where astrocyte conditioned medium was generated in the presence of recombinant IFN-γ. General Linear Modelling containing 1st order interactions p<0.05 was considered statistically significant. IFN-γ: interferon-γ; TNF-α: tumor necrosis factor-α; iRBCs: *P.falciparum*-infected red blood cells; i-lysate: iRBC lysate; nRBC: naive red blood cells; n-lysate: nRBC lysate.

## Discussion

The molecular mechanisms underpinning brain oedema in CM is poorly understood. Histology of brain tissue from CM fatalities shows accumulation of leukocytes and platelets in the brain microvessels throughout the cerebral cortex in areas of iRBC accumulation (73). Similar observations have been made in the cerebral cortex of C57BL/6J mice infected with *Pb*A (73, 74). Vascular activation stemming from BMEC activation is a key event in BBB disruption. BMECs express tight and adherens junction proteins that keeps them tightly adhered to each other minimizing paracellular leak (75). Under steady state conditions, the BBB is maintained with the formation of tight junctions between adjacent BMECs that cover the perivascular space which is populated with astrocytes, pericytes and microglial cells. The BBB-specific traits that BMECs have are, in part, conferred by signals from other cells of the NVU, including glia and pericytes, which are critical for maintaining BBB function (76).

Although evidence exists that glia are activated during *Plasmodium* infections both by the presence of glia-derived molecules in the cerebral spinal fluid of children with CM(77) and in pediatic autposy samples in fatal CM cases (78), very little mechanistic data outlining the contribution of glia to BBB disruption in *Plasmodium* infection exists. Astrocytes and microglia are both activated in ECM (12, 79, 80) providing a tractable model to determine how glia activation may contribute to the breakdown of the BBB during *Plasmodium* infection.

IFN-γ is required to mediate BBB breakdown in ECM (74). The bulk of IFN-γ comes from activated CD4 T cells (16) augmenting cross-presentation of merozoite-derived antigens by BMECs and the trafficking and recognition of parasite antigen by pathogenic CD8 T cells (81). CD8 T cells are critical mediators of BBB dysfunction in ECM (15, 20) and accumulate in the brain during pediatric CM (14, 82). Whilst CD8 T cell-mediated apoptosis of BMECs is not widespread making it unlikely to be the major contributor to vasogenic odema, the down-regulation of BMEC expressed junction proteins is significant.

The data presented here demonstrate that in the ECM model astrocytes have a strong IFN signature (**Fig. 1**). Although this does not completely distinguish between type II interferon signalling initiated by IFN-γ and type 1 IFN sigalling through interferon α/β receptor (IFNAR), *in vitro* data from cultured hCMEC/D3 monolayers indicate that IFN-γ does impact astrocyte-derived molecules that may influence the ability of BMECs to form cohesive monolayers (**Fig. 5**). Surprisingly the trasncriptional response of astrocytes to IFN-γ overcome some of the disruptive BMEC responses to *P. falciparum*-derived molecules *in vitro* suggesting a response that would augment the BBB. This data is in agreement with the transcriptional data obtained from astrocytes in *Pb*A-infected mice (**Fig. 1**) and existing data that removal of IFN-γ signaling on astrocytes / neurons does not appear to improve survival during *Pb*A infection (37). This data also mirrors recently published data suggesting microglia have a protective role against coagulation in a hyperinflammatory model of CM (12, 83). Therefore both astrocytes (**Figs 1 and 5**) and microglia respond to interferon signalling with protective pathways but are unable to overcome the factors leading to downregulation of BMEC junction protein expression in response to *Pb*A infection. It also explains why removal of microglia did not alter the survival of mice with ECM (**Fig. S2**).

Whilst there is a plethora of information on the effects of INF-y signaling on BMECs, the goal of this study was to examine the effects of INF-y signaling on astrocytes. One of the effects of IFN-γ signaling was to reduce the transcription of Wnt7a from astrocytes (**Fig. 1E**) and a decrease in the Wnt7a activity score in BMECs during *Pb*A infection (**Fig. 2C**). Whilst *in vitro*, stimulation of human astrocyte cells with *P. falciparum* lysate resulted in elevated Wnt7a production (**Fig. 3B**) demonstrating that astrocytes can respond directly to products from *Plasmodium* parasites, the addition of recombinant IFN-γ to the cultures abolished Wnt7A production to *P. falciparum* lysed iRBCs (**Fig. 3B**). The role of Wnt7a in BBB integrity appears to highly depend on context. Whilst astrocyte-derived Wnt7A has been shown to be protective in ischemic stroke (24) and supplementation of Wnt7a rescued BBB disruption in a mouse COVID-19 model (25), Wnt7a has also been reeported to disrupt BBB specifically via a decrease in tight junction proteins expression by BMEC mediated by the nuclear translocation of β-catenin (84).

We did not see any improvement in survival during *Pb*A infection in mice where Wnt7A was supplemented at the BBB by an AAV delivery system (42) suggesting that this is not the reason for BBB disruption in this context. Wnt7a activates β-catenin which has been shown to promote endothelial barrier integrity (85, 86) possibly through reinforcing the adhesiveness of inter-BMEC cadherin bonds (87). However in *Plasmodium* infection it has been shown that activation of β-catenin had the opposite effect (32). The pathway to β-catenin activation may be important in this respect with evidence that β-catenin activation via angiotensinogen II could be highly detrimental. BMEC β-catenin activation, or lack of, via Wnt7a activity from astrocytes (**Fig. 2**) may be less important.

In conclusion, glia respond strongly to IFN signaling in *Plasmodium* infection but this appears to induce a protective transcriptional program that ultimately cannot overcome the force of vascular inflammation induced by BMECs responding to parasite antigens. Novel therapeutic strategies for CM may need to overcome the effects of inflammatory pathways in BMECs to facilitate astrocyte-facilitated rescue of BBB integrity.

## Acknowledgements

We thank the Comparative Medicine Center staff at the University of Utah for excellent animal husbandry and the University of Utah Sequencing and Genomics Core Facility in particular Derek Warner and Mike Powers for the generation and sequencing of single nuclear and single cell libraries. We also thank Benoit Vanhollebeke (University of Brussels) for sharing the Wnt7a-expressing AAV. We thank Wendy Zhu and Emily Mayville for technical assistance. This work was supported by a R01NS097819 awarded TJL from the National Institute for Neurological Disorders and Strokes. MR and TJL devised the experiments. MR, NB, RD, KC, LR and TL performed the experiments and analyzed data. JA analyzed the data. MR and TJL drafted the paper. TJL supervised the experiments, analyzed data, and wrote and edited the paper. The funders had no role in study design, data collection and analysis, decision to publish, or preparation of this manuscript.

## Ethics approval statement

All animal procedures were approved by the Institutional Animal Care and Use Committee (IACUC) of University of Utah (Protocol No. 2051). All efforts were made to minimize animal suffering and to reduce the number of animals used, in accordance with the ARRIVE 2.0 guidelines. Euthanasia methods used were consistent with AVMA Guidelines for the Euthanasia of Animals.

---

**Supplemental Figure 1:**
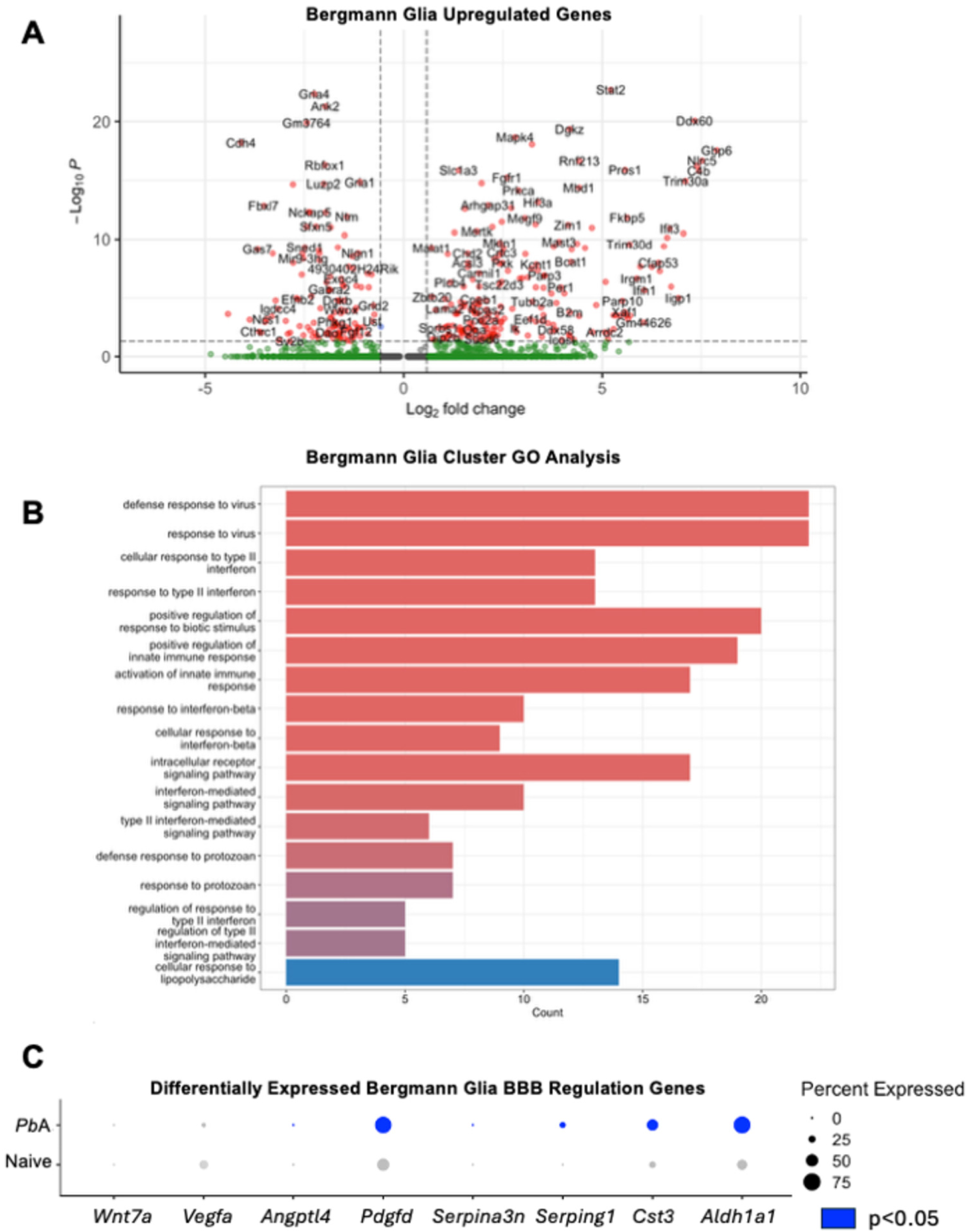
Bergmann glia upregulate interferon response genes and alter BBB regulation genes in response to *Pb*A infection. A volcano plot of differentially expressed genes in the Bergmann glia cluster from the snRNA-sequencing of whole brains of control (naïve) mice compared to mice infected with *Pb*A (**A**). Gene ontology (GO) analysis of the 200 most highly upregulated genes in the Bergmann glia cluster from *Pb*A-infected mice relative to naive mice (**B**). A dot plot of genes that regulate blood brain barrier function that are significantly differentially expressed in the Bergmann glia of *Pb*A*-*infected mice relative to naive mice (**C**). p<0.05 was considered statistically significant.

**Supplemental Figure 2:**
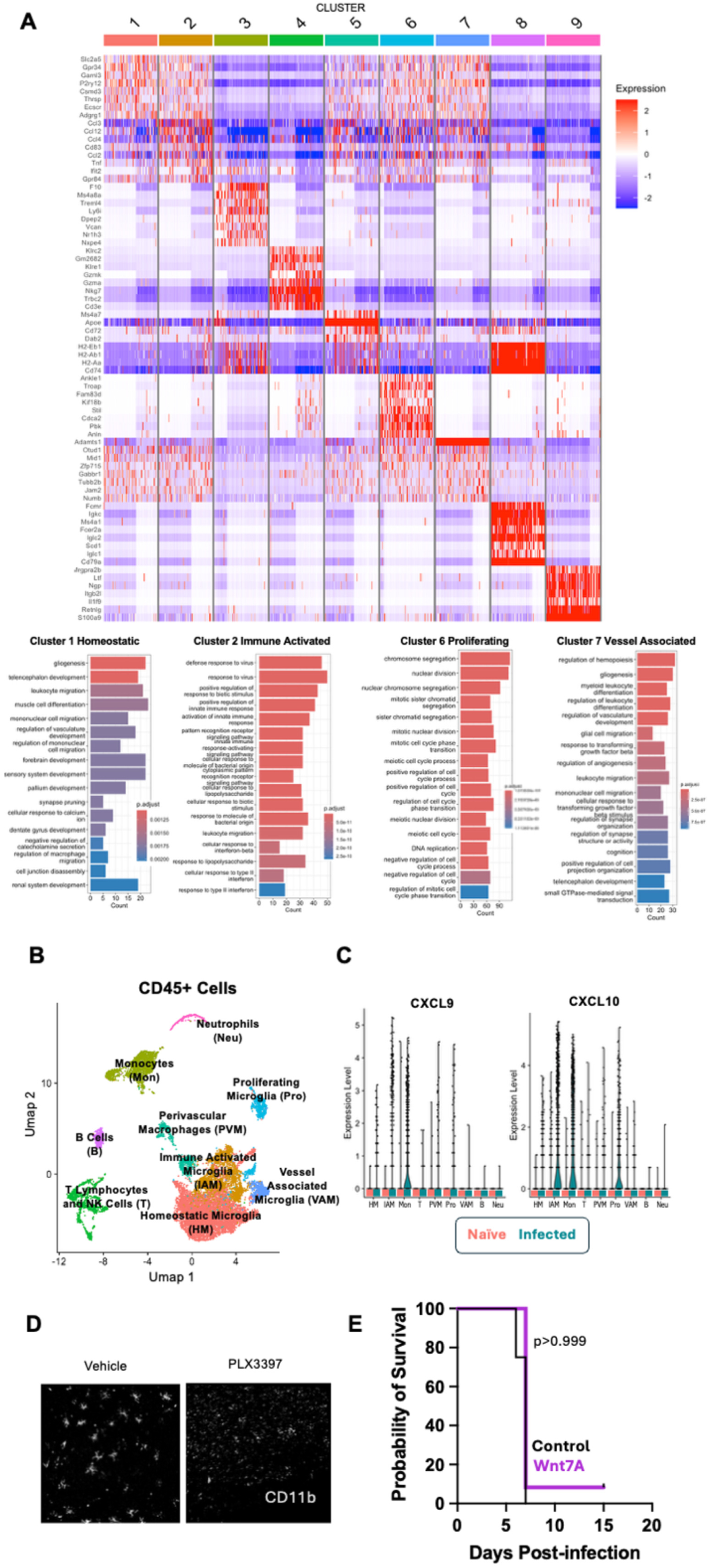
Depletion of microglia does not protect against death in *Pb*A-mediated BBB disruption. Heatmap of top markers of each cluster from single-cell RNA-sequencing of CD45+ cell (naïve) and mice infected with *Pb*A (**A**). The gene ontology (GO) analysis of the 500 most highly upregulated genes in 4 clusters that group microglia is shown for clusters 1, 2, 6 and 7. Of the 9 clusters (**B**), in *Pb*A-infected mice, monocytes (mon) have increased transcription of CXCL9 and CXCL10 whilst Immune Activated microglia (IAM) have increased transcription of CXCL10 and proliferating microglia (pro) have increased transcription of CXCL9 (**C**). HM: homeostatic microglia; T: T cells and natural-killer cells; B: B cells;, Neu: Neutrophils; PVM: peri-vascular macrophages; VAM: vessel associated microglia. Confocal images of brain tissue of mice treated with vehicle control or PLX3397 and stained with CD11b to visualize microglia (**D**). Probability of survival of intact (n=8) or microglia-depleted (n=8) *Pb*A-infected mice (**E**). A difference in survival was tested using a Log-rank Mantel-Cox test. p<0.05 was considered statistically significant in all cases.

**Supplemental Figure 3:**
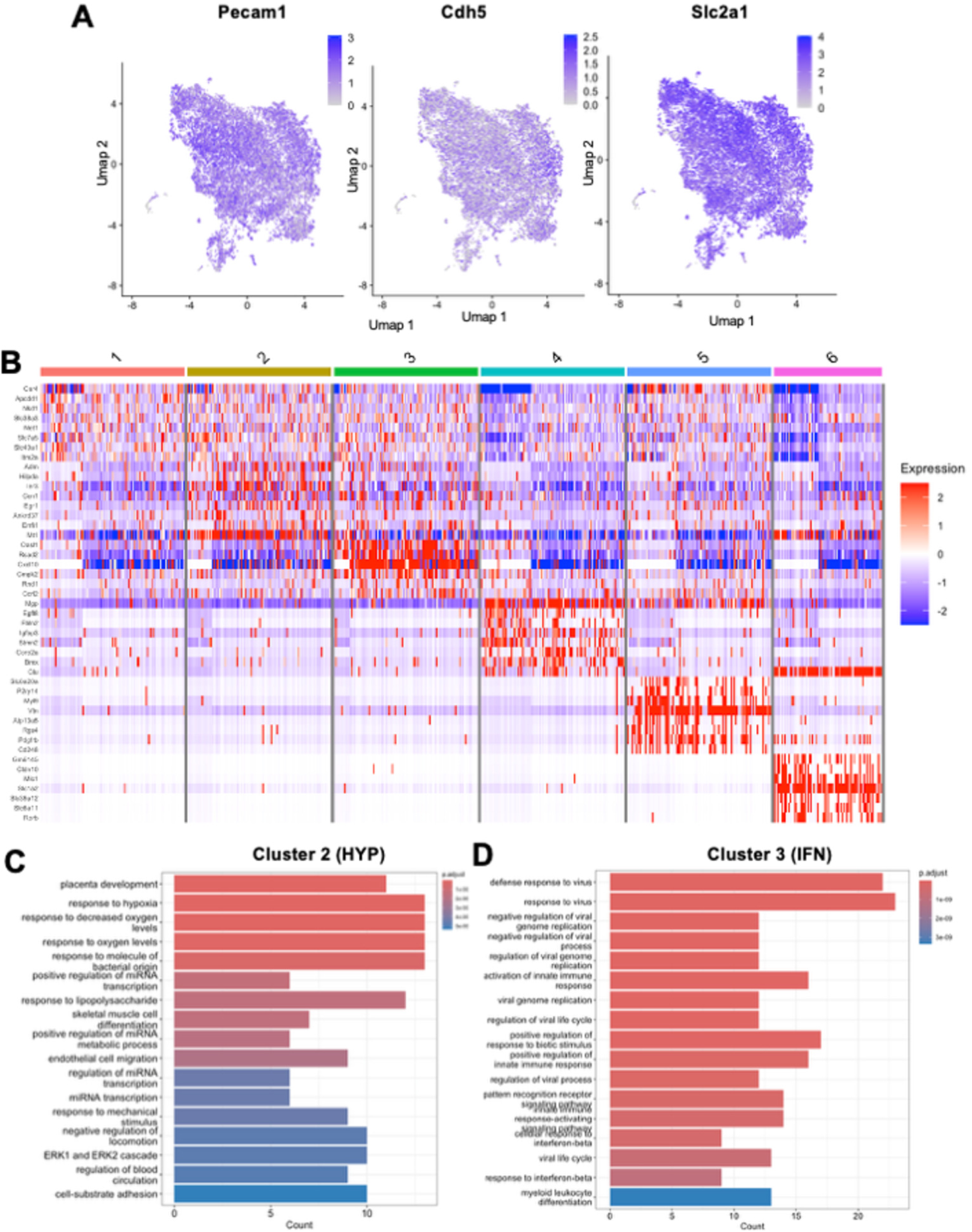
Transcriptional changes in BMECs from *Pb*A-infected mice reflect different activation states. Magnetic isolation of CD45-/CD31+ cell yielded a scRNA sequencing dataset where cells expressed high levels of endothelial markers *Pecam1*, *Cdh5* and *Slc2a1* (**A**). Heatmap of highly expressed markers in the 6 clusters of BMECs generated (**B**). The gene ontology analysis of the 100 most highly upregulated genes in cluster 2 indicative of a hypoxic response state (HYP) (**C**) and cluster 3 with significant interferon signaling pathways activated (IFN) (**D**). p<0.05 was considered statistically significant.

